# *Monodelphis domestica* as a fetal intra-cerebral inoculation model for Zika virus pathogenesis

**DOI:** 10.1101/785220

**Authors:** John M. Thomas, Juan Garcia, Matthew Terry, Ileana Lozano, Susan M. Mahaney, Oscar Quintanilla, Dionn Carlo-Silva, Marisol Morales, John L. VandeBerg

## Abstract

*Monodelphis domestica*, also known as the laboratory opossum, is a marsupial native to South America. At birth, these animals are developmentally equivalent to human embryos at approximately 5 weeks of gestation which, when coupled with other characteristics including the size of the animals, the development of a robust immune system during juvenile development, and the relative ease of experimental manipulation, have made *M. domestica* a valuable model in many areas of biomedical research. However, their suitability as models for infectious diseases, especially diseases caused by viruses such as Zika virus (ZIKV), is currently unknown. Here, we describe the replicative effects of ZIKV using a fetal intra-cerebral model of inoculation. Using immunohistochemistry and in situ hybridization, we found that opossum embryos and fetuses are susceptible to infection by ZIKV administered intra-cerebrally, that the infection persists long term, and that the infection and viral replication consistently results in neural pathology and may occasionally result in global growth restriction. These results demonstrate the utility of *M. domestica* as a new animal model for investigating ZIKV infection *in vivo.* This new model will facilitate further inquiry into viral pathogenesis, particularly for those viruses that are neurotropic, that may require a host with the ability to support sustained viral infection, and/or that may require intra-cerebral inoculations of large numbers of embryos or fetuses.

**AUTHOR SUMMARY:** Here we show that the laboratory opossum (*Monodelphis domestica*) is a valuable new model for studying Zika virus pathogenesis. Newborns are at the developmental stage of 5-week human embryos. Zika virus inoculated on a single occasion into the brains of pups at the human developmental stages of 8-20 weeks post conception replicated in neuronal cells and persisted as a chronic infection until the experimental endpoint at 74-days post infection. In addition, we observed global growth restriction in one of 16 inoculated animals; global growth restriction has been observed in humans and other animal models infected with Zika virus. The results illustrate great potential for this new animal model for high throughput research on the neurological effects of Zika virus infection of embryos and fetuses.

## INTRODUCTION

Zika virus (ZIKV) is a small, enveloped positive-sense RNA virus from the family *Flaviviridae*. Typically transmitted in a zoonotic cycle that alternates between a vertebrate host and an invertebrate vector, ZIKV gained notoriety following the 2015 outbreak in Brazil, which saw a dramatic increase in the number of neurological abnormalities in infants born to ZIKV-infected mothers [1]. Significant increases in Guillain-Barre syndrome and microcephaly during this outbreak were also observed when compared to previous years [2], perhaps fulfilling the theory posited by Hayes when he declared ZIKV to be neurovirulent [3].

Following the initial isolation of ZIKV from the upper canopy of the Ziika Forest in Uganda in 1947 [4], little research into the neuropathology of ZIKV had been carried out prior to the Brazilian epidemic. It is estimated that more than 400 babies were born with microcephaly and other brain abnormalities to ZIKV-infected pregnant women during this outbreak [5] and, subsequently, analysis of fetal tissue collected from ZIKV-infected infants supports a causal relationship between ZIKV and neurological abnormalities, as ZIKV has been detected in brain tissue of microcephalic fetuses, as well as in amniotic fluid of pregnant women [6, 7, 8]. The dramatic increase in the incidence of microcephaly and other fetal abnormalities from the Brazilian outbreak has spurred the development of animal models of infection in order to study the effects of ZIKV replication *in vivo*, with a particular focus on the neurotropism of ZIKV. To date, the principal animal models for assessing ZIKV pathology have been nonhuman primates (NHPs) and transgenic mice; limited studies also have been conducted with chicken embryos [9].

The NHP model is the most relevant in terms of reproducing the pathology *in vivo* compared to what is known about ZIKV-induced pathologies in humans [10, 11, 12, 13]. Macaques have been the NHP model used most frequently, and several studies have demonstrated the advantages of the NHP model by comparison with the mouse model, including similarities to human gestation, ease of studying placental transmission, and robust immune responses as expected in an immuno-competent animal model [10, 14]. In a fetal macaque model, ZIKV elicited severe pathological effects on the central nervous system (CNS) including damage to the axonal and ependymal area, gliosis, and hypoplasia of the cerebral white matter [13]. Other studies using NHPs have shown high viral loads in the sex organs and consistent viral shedding in the oral mucosa, further suggesting that NHPs may be uniquely suited to addressing many questions pertaining to ZIKV pathogenesis *in vivo* [10, 15]. However, the cost associated with the use and maintenance of NHPs precludes large-scale experimentation. This limitation, together with the long duration of time required to investigate effects of ZIKV infection during gestation on development during infancy, adolescence, and into adulthood and old age in NHPs, suggests that additional animal models are required.

Studies conducted with immune-deficient murine models have demonstrated the ability of ZIKV to replicate in neuronal and ocular tissue [16], to delay development and whole-body growth [17], to reduce cortical thickness and cell numbers [17], and to elicit apoptosis in ZIKV-infected neurons [18]. Many of the immune-deficient murine models are based on the abatement of type I interferon responses (A129 mice) or types I and II interferon (AG129 mice) [19]. These animals are highly susceptible to ZIKV infection, maintain a high viral load in the CNS, and demonstrate the ability of ZIKV to infect cells associated with the testes, an observation that is consistent with the findings of sexual transmission of ZIKV from males to females in humans [20, 21]. The transgenic murine models have generated useful data regarding ZIKV pathogenesis; however, because these animals are deficient in cell-mediated immune responses that are often the most effective defense against intracellular pathogens [22], data from transgenic murine models may not be fully representative of the pathology observed in humans.

Other studies have used sub-cutaneous ZIKV infection of immunocompetent 1-day-old C57BL/6 pups (immunocompetent to the limited extent that 1-day-old mouse pups have begun to develop their immune system), which resulted in the development of major brain abnormalities including neuronal cell death, gliosis, and axonal rarefaction [23], all of which are representative of ZIKV replication in human brain tissue [2, 24, 25,]. From a developmental perspective, however, a 1-day-old mouse pup is approximately equivalent to a human fetus at 19-weeks of gestation [26] and, as such, is not suitable for modeling the effects of ZIKV infection of human embryonic and earlier fetal stages. Moreover, C57BL/6 pups inoculated at 3 or 10 days of age did not develop any signs of disease, so the neonatal mouse model is limited to a single stage of fetal development.

In an effort to model ZIKV infection of those stages of human development, Shao et al. [27] performed intra-cerebral inoculations of e14.5-day mouse embryos with ZIKV and allowed them to develop. An e14.5 mouse embryo is developmentally equivalent to a human embryo at 7-8 weeks post-conception [26]. Massive neuronal death occurs in the inoculated embryos and, although it is possible for some of them to survive to birth, the oldest animal reported was 3 days old, suggesting that the infection is lethal within days of birth. Because of the time and effort required for inoculating mouse embryos, this model is not practical for high throughput experiments that are required for modeling the various potential outcomes of human embryo infection with ZIKV. Moreover, since the infection is lethal in this model, it is not possible to use it to investigate long-term sequelae of ZIKV infection at the embryonic stage.

Because all of the existing animal models of ZIKV-induced pathogenesis have significant limitations, we explored the potential of a marsupial model to circumvent those limitations. The gray short-tailed opossum, *Monodelphis domestica*, is native to Brazil and surrounding countries. The laboratory genetic stocks and inbred strains of this species are collectively referred to as the laboratory opossum [28]. Laboratory opossums are widely used as models in many fields of biomedical research [29], and they possess some characteristics that render this model suitable, and in some respects, unique, for studying the pathogenesis of ZIKV *in vivo*. First, the animals are small (80-140g as adults), but several times the size of a mouse, facilitating some experimental procedures by comparison with mice, such as serial collections of substantial quantities of blood. Second, they are highly fecund, and easy to manipulate; and they can be produced and maintained cost-effectively. Third, at birth, *M. domestica* are developmentally equivalent to a human embryo at approximately 5 weeks of gestation [26], and they complete embryonic and most of fetal development while attached to the mother’s nipples over a 2-week period [30]. Fourth, female *M. domestica* do not have pouches, so the pups can easily be experimentally manipulated while they are attached to the nipples, and they have a high rate of survival post-manipulation. Fifth, while the immune system is undeveloped at birth, *M. domestica* develop a fully intact immune system as they develop beyond the fetal stage. Last, as is also true for the immune-deficient murine models, but not for the in utero murine model or the NHP model, large numbers of *M. domestica* can be used economically, enabling robust statistical analysis for between-group comparisons, as well as robust assessment of within-group variations in outcome of ZIKV infection.

We emphasize the importance of being able to assess within-group variation in large numbers of animals inoculated with ZIKV, for the purpose of modeling the major variations in pathological outcome of human infection with ZIKV. For example: 1) growth retardation and microcephaly are uncommon outcomes of ZIKV infection of human embryos and fetuses [24]; eye pathologies occur in only a minority of children who were infected in utero and in only a small proportion of children and adults who become infected with ZIKV [6, 31]; Guillain-Barre syndrome is caused by ZIKV infection in only a minority of people [32].

The purpose of this study was to assess the utility of *M. domestica* as an intra-cerebral model for ZIKV neuropathogenesis by determining if ZIKV can replicate and persist in the brains of young pups and, if so, to determine the nature and extent of the neuropathological consequences by comparison with those observed in humans and other animal models.

We point out that the experiment reported here is not intended to model the complex biological processes that lead to infection of brains of human embryos and fetuses with ZIKV, typically beginning with the bite of a mosquito, replication in the mother, trans-placental transfer to the embryo or fetus, replication in the embryo or fetus, followed by entry into the brain and replication in the brain. Rather, our model obviates all of the variables and mechanistic complexities that exist between the time of initial infection of the mother and entry of the virus into the brain of the embryo or fetus. Via the use of this unique model, we can conduct high throughput experiments to investigate short-term and long-term pathological effects of variation in the number of PFU that enter the brain, the exact developmental time point at which they enter the brain, and the genetic make-up of different ZIKV strains, in the absence of the many confounding variables that exist in models of trans-placental infection of embryos and fetuses.

## METHODS

### Animals

The laboratory opossums used in this study were produced in the breeding colony maintained at The University of Texas Rio Grande Valley and maintained under standard conditions [28].

### Ethics Statement

All animal work described herein was subject to review and approval by the UTRGV Institutional Animal Care and Use Committee (IACUC), as well as oversight provided by the UTRGV Department of Laboratory Animal Resources (LAR). LAR maintains compliance with the National Institutes of Health Office of Laboratory Animal Welfare (NIH OLAW) Public Health Service (PHS) Policy on Humane Care and Use of Laboratory Animals; PHS Assurance number A4730-01, and the United States Department of Agriculture (USDA); USDA Assurance number 74-R-0216. The animal protocol for this work was approved, and conducted under the IACUC protocol of Dr. John Thomas (#2016-005-IACUC).

### Susceptibility of *M. domestica* pups to ZIKV infection

In the first experiment, *M. domestica* pups were inoculated with 5,000 PFU of ZIKV PRVABC59 intra-cerebrally. Two litters at 2 and 3 days of age, respectively, were used, hereafter referred to as Group 1 and Group 2. Each group contained three animals. At 19 days post-inoculation, the animals were euthanized, and whole brains were collected, weighed, and homogenized for virus titration.

### Developmental effects of intracerebral inoculation

Following the initial confirmation that *M. domestica* pups could be infected with ZIKV via the intra-cerebral route, in the second experiment we examined the effects of ZIKV infection on postnatal development in the laboratory opossum model. *M. domestica* pups ranging in age from 4-20 days (equivalent in human development to 8-20 weeks post conception) were inoculated intra-cerebrally with 5,000 PFU of ZIKV as described above. Control animals, ranging in age from 2-9 days, were inoculated with PBS. Seventy-two days after the inoculations, the animals were euthanized, weighed, and measured; and brain tissue was collected for analysis by immunohistochemistry and in situ hybridization.

### Cells and viruses

ZIKV isolate PRVABC59 (a gift from Dr. Kenneth Plante at the WRCEVA repository at UTMB) was used for the inoculations. Vero cells (CCL-81; ATCC, USA) were used for virus titration, and C6/36 cells (CRL-1660; ATCC, USA) derived from *Aedes albopictus* were used to amplify lyophilized virus for scale-up. Virus generated from the initial reconstituted lyophilized stock was passaged once in C6/36 cells, and the resulting supernatant was clarified and purified over a sucrose cushion. Virus supernatants were quantified in duplicate by plaque assay, as described previously [33]. Aliquots were stored at −80°C for further use.

### Tissue fixation and sectioning

Dissected tissue was fixed in sterile PBS (Gibco, USA) + 4% formaldehyde solution and stored at room temperature. Fixative was then cleared from tissue by performing three quick washes in sterile PBS followed by three 10-min washes in sterile PBS. Next, the tissue was washed 1X for 5 min in a 25% methanol:PBS solution; washed 1X for 5 min in 50 % methanol:PBS solution; and finally washed 3X for 5 min in 100% methanol. Tissue was stored at −20°C until needed. Tissue was rehydrated by washing 1X for 5 min in 50% sterile methanol:PBS; washed 1X for 5 min in 75% methanol:PBS, and then washed 3X for 5 min in sterile PBS. Tissue was incubated for 30 min in 33% OCT mounting media: sterile PBS, 30 min in 66% OCT: sterile PBS, 1-4 hours in 100% OCT. Tissue was mounted in OCT and cooled to −20°C for sectioning by a cryostat (Leica Biosystems, USA). Sections of 10 – 20 µm were mounted onto Frost + microscope slides and stored at −20°C.

### Antibody staining

Mounted sections of tissue were incubated in PBTB (sterile PBS + .01% Tween20 + 0.2% BSA) for 1 hour followed by incubation in 1:500 dilution of primary antibody (Arigo Biolaboratories, Taiwan) for either 1 hour at room temperature or overnight at 4°C. Primary antibody was removed by washing 3X quickly, then 3X for 10 min each in PBTB. Tissue was then incubated in 1:200 dilution of AlexaFluor (546 or 647 – Thermo Fisher Scientific, USA) conjugated secondary antibody in PBTB for 1 hour. Secondary antibody was removed in the same manner as primary antibody, except that DAPI (Thermo Fisher Scientific, USA), and AlexaFluor 488 (Thermo Fisher Scientific, USA) conjugated phalloidin was included in the first 10-min wash. Tissue was imaged using an Olympus FV10i confocal microscope.

### RNA probe preparation

Target genes were amplified using standard PCR and then cloned into PCRII^®^ Expression vector (Invitrogen, USA) as per the manufacturer’s instructions. Cloned products were verified via DNA sequencing and then were linearized by a second PCR using M13F and R primers. After standard PCR cleanup, the linearized gene was quantified and then normalized to 100 ng/µl. Digoxigenin (DIG) tagged probes were made using the SP6 and T7 promoters to make either sense or antisense probes in separate reactions using the following mix: 200 ng linearized cloned PCR product, 2 µl 10X transcription buffer, 1 µl of 0.1M DTT (0.02 M DTT for SP6 reaction), 2 µl of DIG labelled ribonucleotides, 1 µl of RNase inhibitor, 1 µl of either SP6 or T7 polymerase (New England Biolabs, USA). SP6 reactions were incubated at 40°C for 2 hours, and T7 reactions were incubated at 37°C for 1 hour. Successful probe synthesis was confirmed via standard gel electrophoresis, and probes were cleaned using ethanol precipitation and re-suspended in 50 µl of DEPC H_2_O, quantified via spectrophotometry, and stored at −80°C until needed.

### *In situ* hybridization

Mounted sections of tissue were incubated in RNase free PBST (sterile PBS + .01% Tween20) for 5 min, 50% PBST: Hybridization buffer (50% formamide, 5X SSC, 100 µg/ml salmon sperm, 0.1% Tween20, 100 µg/ml Heparin) for five min, then in hybridization buffer for 5 min. New hybridization buffer was placed on the sections, and pre-hybridization was performed in a small, airtight container at 56°C for 2 hours. The working probe solution was prepared by adding ∼200 ng of probe to 100 µl of hybridization buffer and then incubating at 90°C for 5 min, after which the incubation tubes were placed on ice. The working probe solution was then applied to tissue sections and incubated in an airtight container at 56°C for 16 – 24 hours. Probes were washed from the sections using a variety of wash times and numbers, with all washes using hybridization buffer warmed to 56°C and all washes conducted at 56°C. The following protocol was used to minimize non-specific signal: eight washes of 15 min each, followed by four washes of 30 min each. The slides were then cooled to room temperature and washed 1X for 5 min in 50% PBTB: hybridization buffer, 3X for 5 min each in PBTB, 1X for 1 hour in PBTB (to block non-specific protein binding). Slides were then incubated either for 1 hour at room temperature or overnight at 4°C in 1:100 dilution of an HRP conjugated anti-DIG antibody and PBTB. This antibody was removed by three quick washes, and then three 5-min washes in PBTB. Slides were washed in 1X Tyramide buffer for 5 min. Fluorescent labelling was performed using the AlexaFluor Superboost^®^ tyramide signal amplification kit (Thermo Fisher Scientific, USA) following the manufacturer’s instructions and using either the 546 or 647 markers. After the tyramide reaction was stopped, excess reagent was removed by washing 3X quickly, then 3X for 10 min each in PBTB. DAPI and AlexaFluor 488 conjugated phalloidin were included in the first long wash to label nuclei and cytoskeletal elements, respectively. Tissue was imaged using an Olympus FV10i confocal microscope.

### Pathology and NS1 Scoring of Tissues

Brain slices from all animals were prepared, stained, and visualized for detection of NS1 as described above. Tissues were then scored based upon the pathology of the tissue, as well as for expression of NS1. Brain pathology was scored subjectively on a scale of 0-3: 0, normal; 1, mild pathology; 2, moderate pathology; 3, extreme pathology. Brain NS1 levels (extent of fluorescent signal) were scored similarly, using the nuclei visible within the field of view at 60x: 0, none; 1, minimal; 2, moderate; 3, extreme. Images from the PBS control animals were used as an example of normal, uninfected tissue, and established a baseline representation score of 0 (normal morphology; no NS1 signal).

## RESULTS

### Susceptibility of *Monodelphis domestica* to ZIKV infection

Viral replication was detected in two of the three animals from Group 1 (2 days old at the time of inoculation), and one of the three animals from Group 2 (3 days old at the time of inoculation). The average titer was 1.8 × 10^4^ PFU/g of brain tissue from the two 2-day old pups in which virus was detected, while the titer from the single 3-day old pup that was infected was 4.3 × 10^4^ PFU/g of brain tissue (Table 1).

**Table 1.**
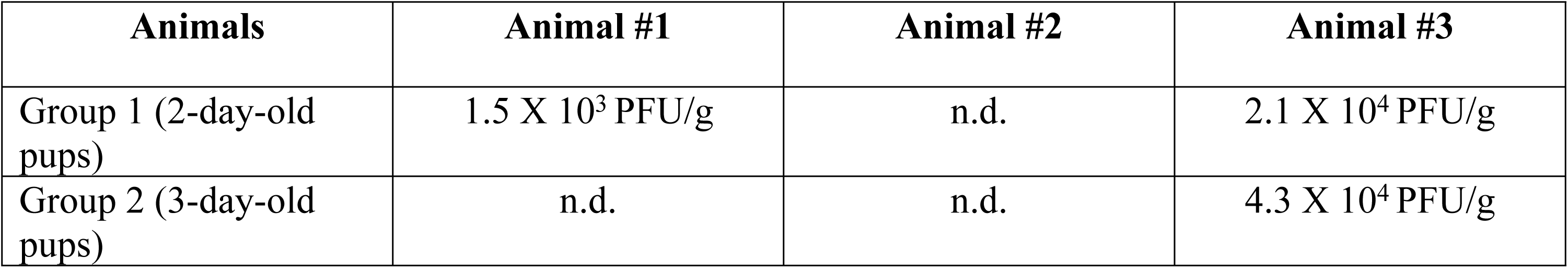
ZIKV replication following intra-cerebral inoculation of *M. domestica*. Three pups in each of two litters of *M. domestica* were inoculated intra-cerebrally with 5,000 PFU of ZIKV PRVABC59. At 19 days and 23 days post-infection, respectively, animals in the two groups were euthanized, and whole brain was collected, weighed, and homogenized for virus titration in Vero cells. Results are the means of duplicate samples. n.d. = no titer detected.

### Developmental effects of intra-cerebral inoculation

One animal (O9355) among the five littermates that were inoculated at 6 days of age and euthanized at 80 days of age had much lower values for body weight, body length, and head length and width, compared to those of its littermates (Fig. 1; Fig. 2). None of the other 10 ZIKV-inoculated animals exhibited growth abnormalities. During 40 years of producing nearly 150,000 laboratory opossums that were not inoculated with ZIKV, we have not observed another animal with such severe growth restriction, although runts are produced on rare occasions. Also, none of the 10 PBS-inoculated animals exhibited growth restriction.

**Figure 1.**
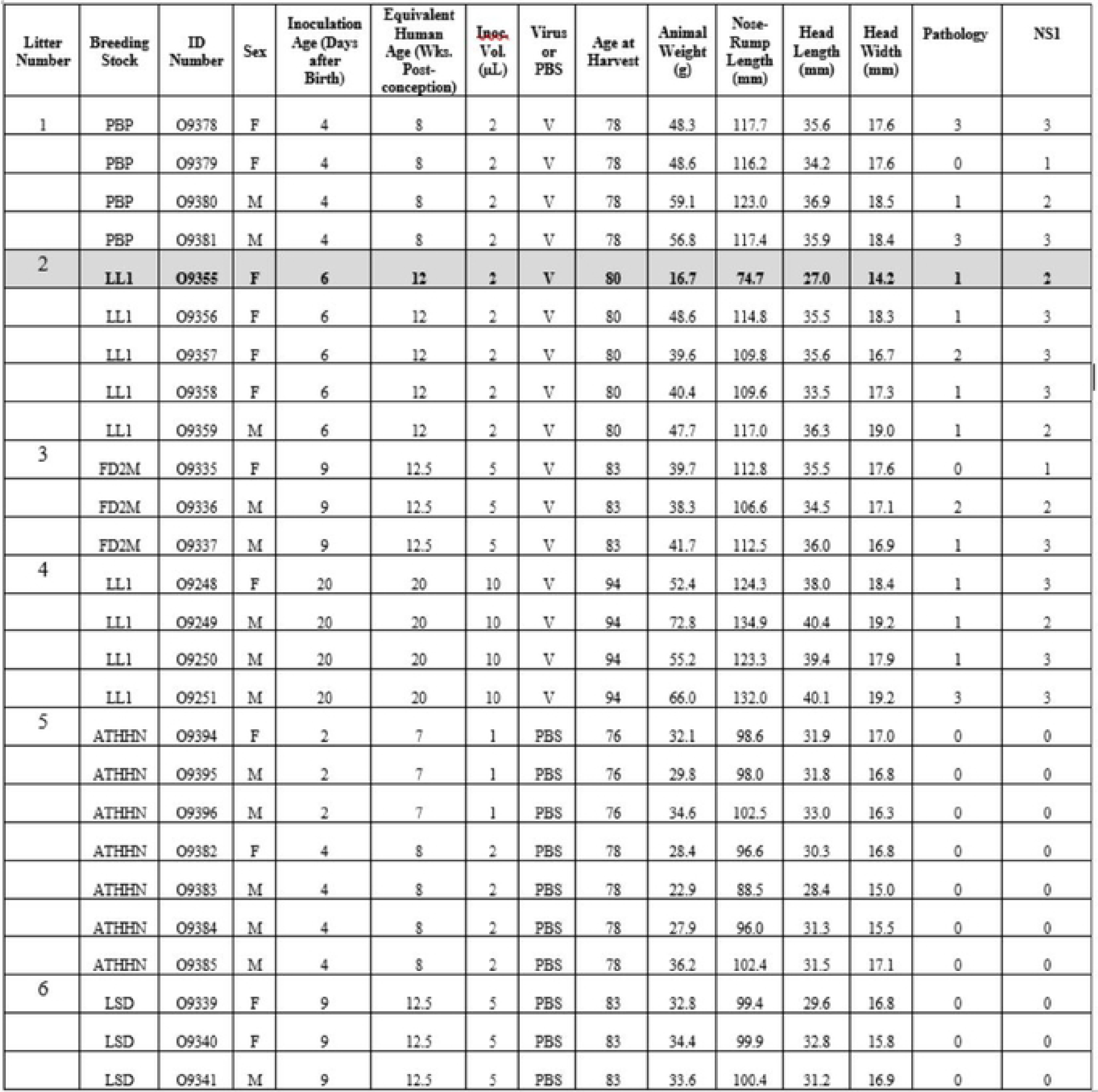
Weights, anatomic measurements, and brain scores from ZIKV-inoculated and PBS-inoculated laboratory opossum pups. Note that weights and measurements among litters are not comparable, because of different ages of harvest and differences among breeding stocks in growth rates. Brain pathology was scored subjectively on a scale of 0-3: 0, normal; 1, mild pathology; 2, moderate pathology; 3, extreme pathology (spongiform-like appearance). Brain NS1 levels (extent of fluorescent signal) were scored similarly: 0, none; 1, minimal; 2, moderate; 3, extreme.

**Figure 2.**
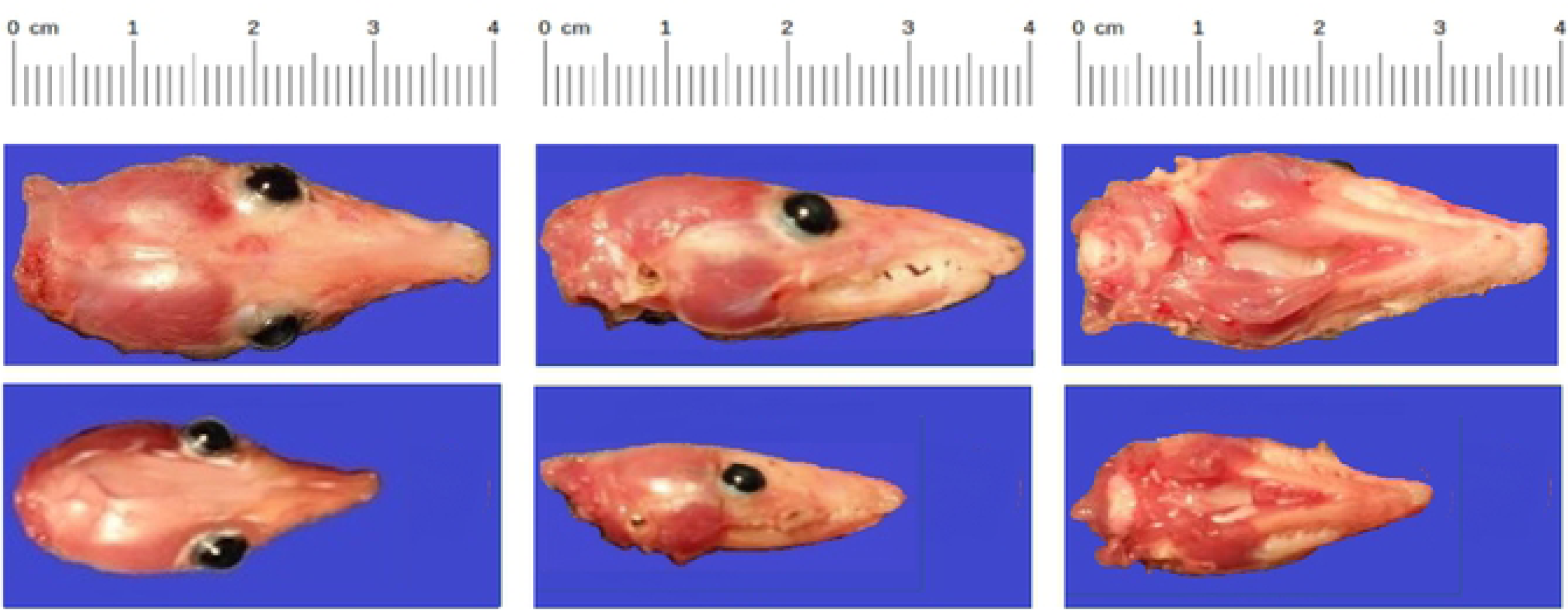
**Heads of normal (top) and growth restricted (bottom) *M. domestica* littermates** at 80 days of age. 6-day-old *M. domestica* pups from a single litter were inoculated with 5,000 PFU of ZIKV PRVABC59 intra-cerebrally. At 74 days post-infection (80 days of age), the animals were euthanized, and photographs and measurements (Fig. 1) of the heads were taken.

### Presence of viral protein and RNA in brains infected with ZIKV

The brains of the pups were fixed, sectioned, and stained for the presence of ZIKV NS1 protein. Immunofluorescence microscopy showed that, in brains collected from all 16 ZIKV-inoculated animals, anti-ZIKV monoclonal antibody directed against NS1 bound specifically to neuronal cells, indicating that the brains were infected with ZIKV (Fig. 3a, 3b). The number of cells visibly expressing NS1 was evaluated and scored based upon the number of nuclei displaying a characteristic punctate staining pattern that we observed in all infected neural tissue samples (Fig. 1). All infected animals showed the presence of NS1 within the brain sections, and the amount of signal appeared to correlate to the pathology of the tissue. Animals that showed the most severe pathology also had the highest number of NS1-fluorescing nuclei, while the samples with more moderate and low pathology scores had low to moderate levels of NS1 expression (Fig. 1).

**Figure 3.**
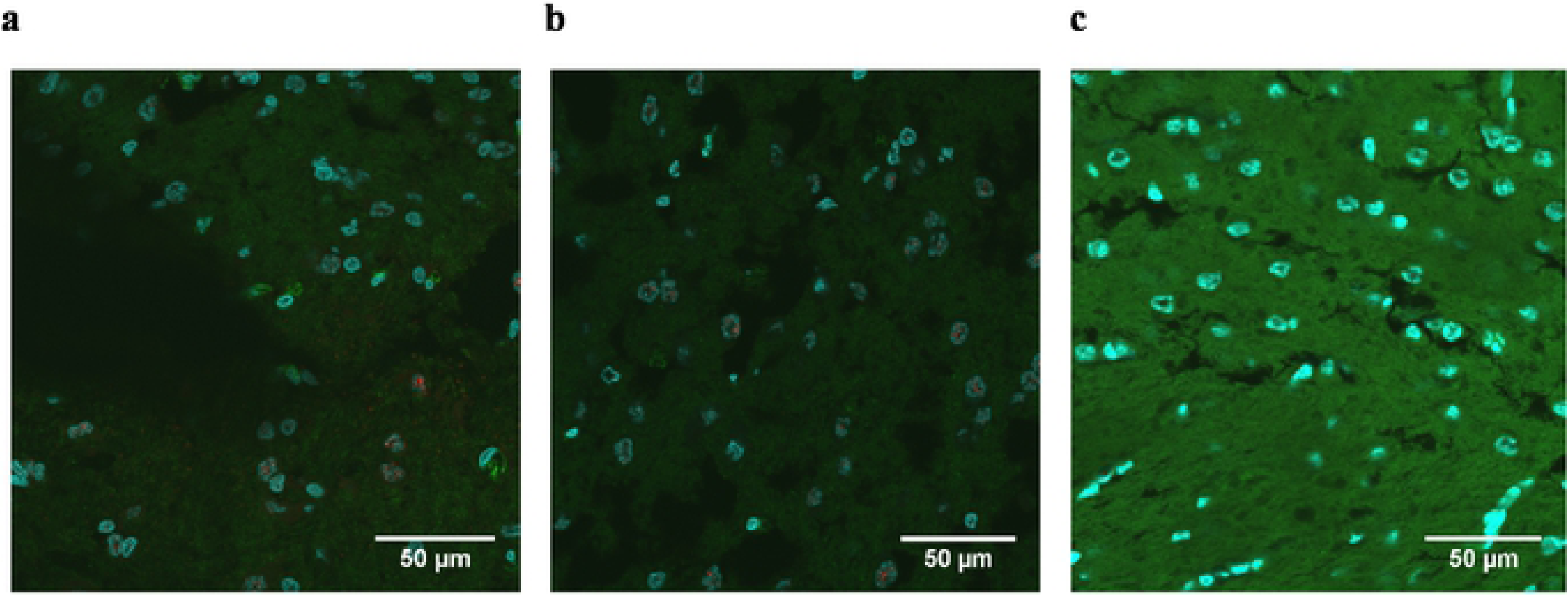
Immunohistochemical detection of ZIKV. **(a)** Immunofluorescence staining of transverse section of cerebellum from infected growth-restricted pup (O9355) at 60x with anti-ZIKV NS1 monoclonal antibody (red). Cytoskeleton is stained green; nuclei are blue. **(b)** Cerebellum section from an infected littermate (O9357). (**c**) Cerebellum section from a mock-infected animal (O9341) at 60x.

Spearman’s correlation coefficient between these semi-quantitative measures of brain pathology and NS1 signal is 0.59 (P = 0.008 for a 1-tailed test of the null hypothesis that there is not a positive correlation between the two measures). Brain tissue sections from each of the 10 animals inoculated with PBS exhibited no evidence of ZIKV (Fig. 3c). To further confirm the presence of ZIKV replication, an *in-situ* hybridization assay was conducted on brain sections of O9355 to detect ZIKV vRNA using NS5 as a target gene. The results showed a strong signal for ZIKV NS5 RNA in the cerebellum (Fig. 4).

**Figure 4.**
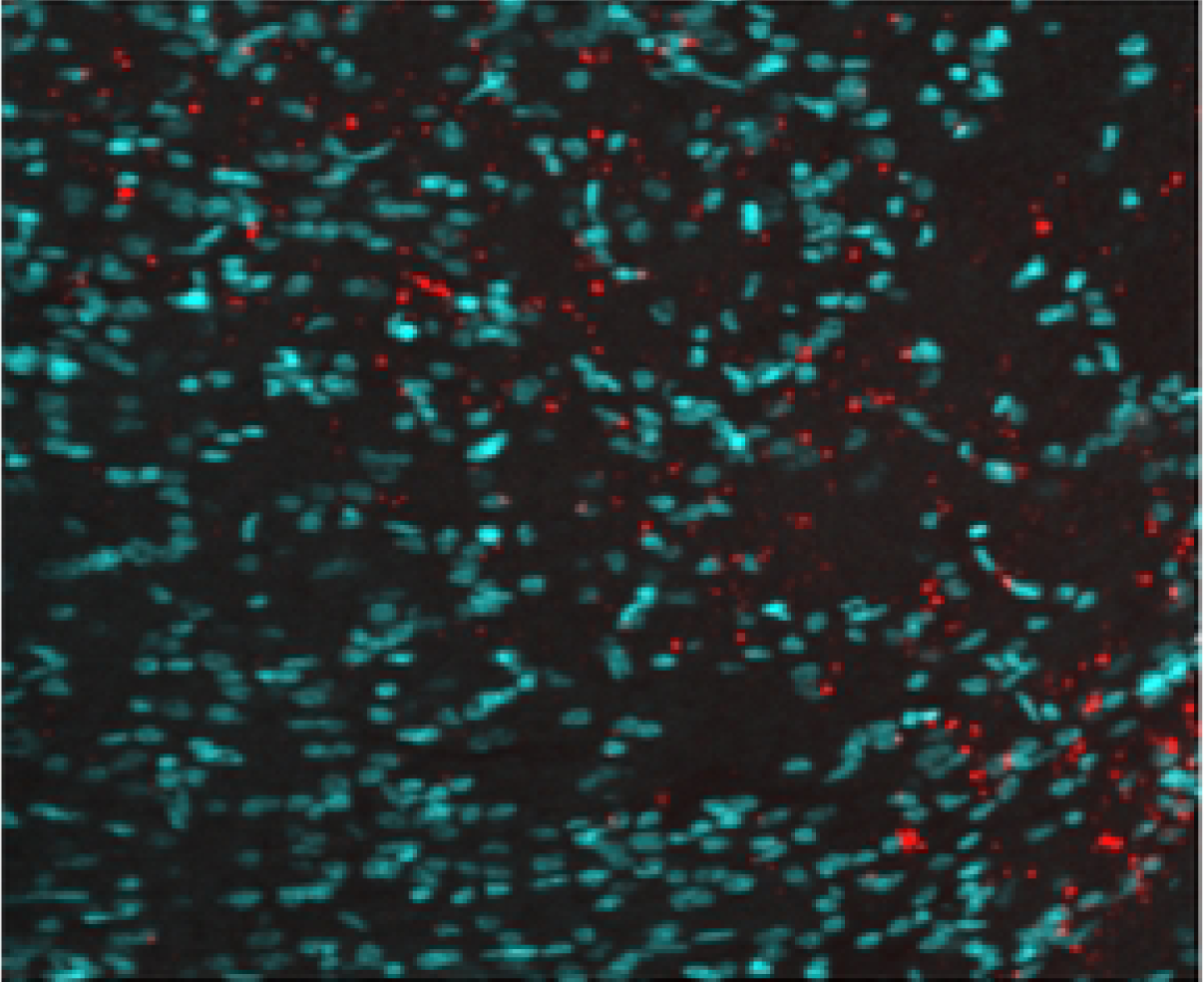
*In-situ* hybridization detection of ZIKV RNA. Immunofluorescence staining of transverse section of cerebellum from infected growth-restricted pup (O9355) at 60x showing the presence of ZIKV NS5 mRNA (red): nuclei are blue.

### Pathological consequences of ZIKV infection in the brain

In addition to demonstrating the presence of ZIKV RNA and protein in the brains of infected pups, images of the fixed neural tissues collected from infected opossum pups were also evaluated based upon the observable cell morphology and scored for severity of disease (Fig. 1). Most of the brain slices showed either a mild or moderate pathology; however, three samples displayed a discontiguous, spongiform-like pathology with large gaps between cells (i.e., a dramatic reduction in density of cells), along with large clumps of DAPI-stained (blue) DNA, which apparently had been released from cells as they died and which had aggregated into large extra-cellular clumps (Fig. 5a). Further examination of brains with this spongiform morphology showed the presence of high levels of ZIKV NS1 protein (Fig.5b). The cerebellum slices from the PBS-inoculated control animals had a uniform, contiguous appearance with little to no gaps between cells, no apparent destruction or cell death, and no extracellular DNA (Fig. 5c-d).

**Figure 5.**
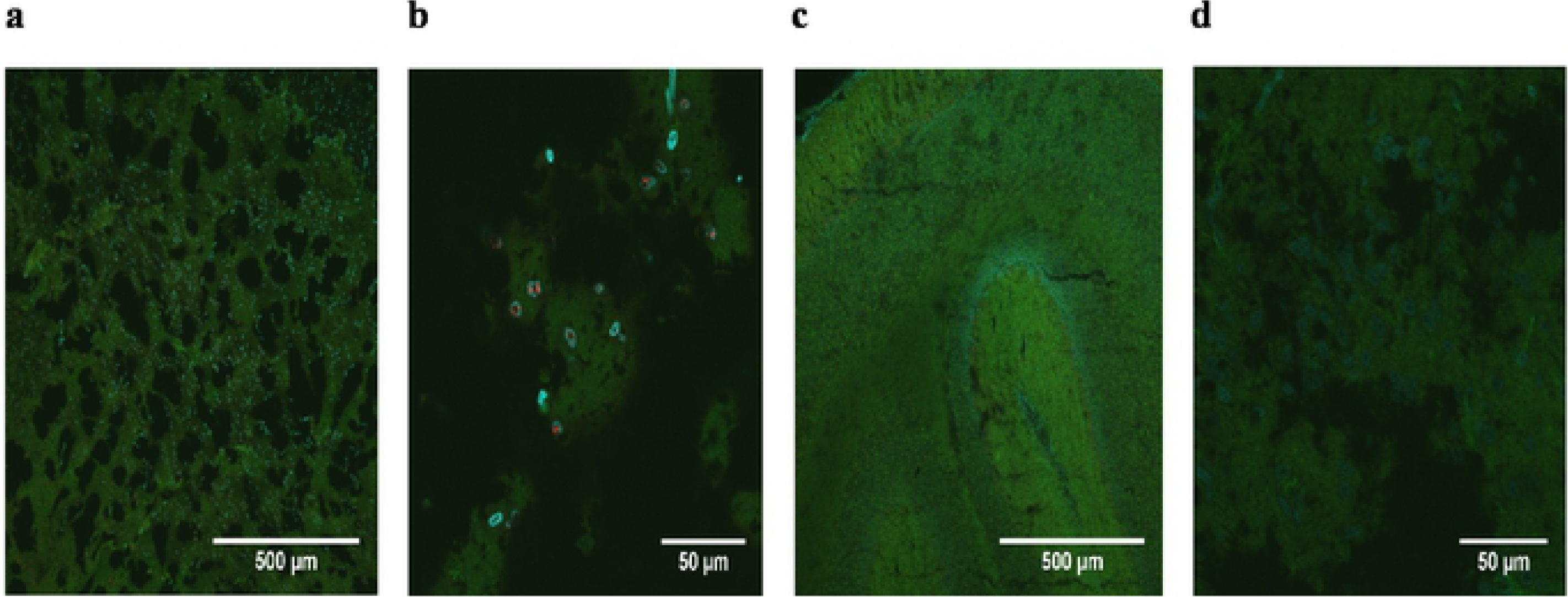
Spongiform morphology induced by ZIKV infection. **(a)** Immunofluorescence staining of transverse section of cerebellum from an infected pup (O9251) at 10x showing a spongiform-like pathology in the presence of ZIKV NS1 protein (red). Cytoskeleton is stained green; nuclei are blue. **(b)** Cerebellum from the same animal at 60x. (**c**) Cerebellum from a mock-infected animal (09339) at 10x. (**d**) Cerebellum from the same mock-infected animal (09339) at 60x.

## DISCUSSION

Two critical questions that pertain to development of a new animal model of infection are 1) is the target host organism susceptible to infection and replication of the pathogen, and 2) does the pathology presented in the animal model accurately reproduce at least some of the clinical findings seen in cases of human infection? As mentioned above, flaviviruses such as dengue virus (DENV) and ZIKV grow poorly, or not at all, in non-primate animals with intact immune systems [34]. Indeed, this lack of susceptibility to viral infection has led to the development and use of immunocompromised transgenic mice and chicken embryos as potential models for ZIKV infection [9]. While these models have demonstrated some utility within the context of understanding ZIKV biology, abatement of the primary immune responses directed against viruses for the purpose of establishing infection may hinder the interpretation of results within the context of relevance to human subjects. Normal, wild-type immunocompetent 1-day-old mice (to the limited extent that mice have a competent immune system at that early age) have been used to model aspects of ZIKV replication and pathology [23]; however, 1-day-old mice correlate with a human fetus at 19 weeks of gestation (20 weeks is mid-gestation) [26]. In contrast, a newborn *M. domestica* pup developmentally correlates to a human embryo at 5 weeks of gestation, thus allowing for ZIKV infection in newborn opossum pups to better replicate the pathology in the developing human embryo during the time when cellular differentiation in critical areas such as the brain are at an early stage. Therefore, the laboratory opossum model, in which ZIKV infection at the embryo or early fetal stage can persist long term, and which can be used in large numbers experimentally, is capable of contributing to our understanding of ZIKV-induced pathologies similar to those that are initiated in humans at early developmental stages.

Due to the potential severe consequences of ZIKV replication in human brains, and its causal association with neurological diseases such as microcephaly, encephalitis, and Guillain-Barre syndrome [25, 35], the ability of *M. domestica* pups to support viral replication in neuronal tissue is an important first step in the validation of the ZIKV laboratory opossum model. Viral amplification of ZIKV and its long-term persistence following a single intra-cerebral inoculation of 4-to 20-day-old animals demonstrated that: 1) brain cells of this species are permissive to ZIKV replication and, 2) this replication ultimately results in cell death and tissue degradation. The ability of fetal *M. domestica* to support viral infection via the intra-cerebral route is not surprising, as fetal mouse brains also support ZIKV infection [23, 27]. However, the long-term survival and continued replication of ZIKV in the brains of *M. domestica* inoculated as embryos or fetuses was a profoundly different outcome from that which occurs with mice. Analysis of the fixed neural tissue showed the presence of ZIKV NS1 protein diffused throughout the tissue, as well as massive cellular death in the brain compared to age-matched sham-inoculated control animals. The presence of NS1 and its distribution across the cerebellum shows that ZIKV replication was persistent for 74 days beyond the inoculation of the virus and suggests that neuronal cells in varied states of differentiation were exposed to ZIKV. This could explain the global growth restriction we observed for one animal and would be consistent with the selective neuronal vulnerability to ZIKV observed in humans [7]. The NS1 protein of ZIKV is a homodimer that, based upon predicted and known NS1 genetic sequences for other flaviviruses, interacts with a variety of host immune factors [36, 37] and is the major antigenic marker of flavivirus infection [38]. The intracellular form of NS1 is central to viral replication, whereas secreted and membrane bound NS1 have been implicated in the excitation of the immune response [38]. The detection of high levels of ZIKV NS5 RNA 74 days post-infection in the one animal (O9355) examined by in situ hybridization confirms that the presence of NS1 detected by immunohistochemistry in the brains of all ZIKV-inoculated pups reflects persistent, active infections in the cerebellum at the time of euthanasia.

A critical finding of our study was the correlation between pathology and level of NS1 signal in the brains of ZIKV-inoculated pups. We consider the correlation of 0.59 (P = 0.008) to be exceptionally high, given that these continuously distributed phenotypes were each subdivided into discrete categories (four for pathology, ranging from 0 to 3; and three for NS1, ranging from 1 to 3, since no ZIKV-inoculated pups scored 0 for presence of NS1). The only three animals that had brains with a spongiform-like pathology (score of 3) all also had the highest score (i.e., 3) for NS1. The only two ZIKV-inoculated pups that had no observable brain pathology (score of 0) had the lowest score for NS1 (i.e., 1) for animals that were inoculated with ZIKV. These results establish that 1) some animals in which ZIKV has been present in their brains since the embryo or fetal stage exhibit no obvious brain pathology, 2) there is variation among littermates in extent of NS1 detected in the brains and consequent extent of pathology, 3) pups as young as 4 days of age and pups as old as 20 days of age at the time of inoculation can develop severe (spongiform-like) brain pathology. Those ages are developmentally equivalent to humans at 8 weeks post conception to 20 weeks post conception (i.e., mid gestation) [26].

Another critical finding was the reduction in overall body size of one infected animal compared to its ZIKV-infected littermates or mock-infected control animals. Surprisingly, the brain of the affected animal (O9355) exhibited only mild pathology (score of 1, see Figure 1), suggesting that growth restriction may not be correlated with overall brain pathology, but rather might be a consequence of a localized perturbation in brain development caused by ZIKV infection. While the sample size was small, the physical measurement data suggest that infection of *M. domestica* with ZIKV at the embryonic stage of development can occasionally result in severe growth restriction. Infection of immunocompetent mouse embryos also can result in growth restriction, and it has been suggested that infection of embryonic mouse brain by ZIKV causes an immune response that disrupts neurovascular development [27]. While initial reports from the WHO and CDC originally highlighted microcephaly as the major concern with vertical transmission of ZIKV infection in pregnancy, more recent studies refer to Congenital Zika Syndrome (CZS), of which microcephaly is one severe manifestation of infection. ZIKV infection of a single pregnant pigtail macaque resulted in several sequelae in the fetus reminiscent of CZS in humans, including restricted fetal brain growth and the presence of viral RNA in the brain [14]. Additionally, infection of pregnant rhesus macaques similarly demonstrated evidence of disrupted fetal growth, prolonged maternal viremia, and inflammation at the maternal-fetal interface, including mild decidual perivascular inflammation (not unusual in human decidua) and placental acute chorioamnionitis [39]. Therefore, evaluation of the opossum model using the expanded criteria of CZS (as has been suggested by others to include, but not be limited to, microcephaly), may allow for a more comprehensive understanding of the neurotropism of ZIKV in the fetal brain. In the opossum model, we observed several manifestations of CZS to include: 1) overall reduction in total body size and weight of one animal; 2) the presence of ZIKV NS1 protein as well as NS5 vRNA in the brains of infected pups; and 3) reductions in total number of glial cells, gliosis, hypoplasia, and cellular damage. As discussed above, we also observed a spongiform-like pathology in three infected animals.

While it is unknown what the long-term sequelae of CZS would be in the opossum model, studies are underway to evaluate the long-term impact of ZIKV infection on growth, development, mental and physical capabilities, and behavior. Indeed, it has been recently shown that postnatal infection of ZIKV resulted in sustained structural and functional alterations in an infant macaque model [11]. This result suggests that ZIKV infection can have deleterious developmental implications that go far beyond the ‘classical’ definition of ZIKV neuropathology in relation to the size and structure of the brain. Indeed, we have demonstrated that subcutaneous, intramuscular, or intraperitoneal inoculation of ZIKV into juvenile laboratory opossums with intact immune systems can result in chronic infection, viral dissemination to many organs including brain and reproductive organs, and anatomic and histological abnormalities (unpublished data).

And, as shown recently, some of the symptoms described in the transgenic murine fetal models do not result in development of microcephaly, suggesting that there may be other factors that influence neurological outcomes in transgenic murine models. For example, studies using pregnant C57BL/6 (Ifnar1-/-) dams infected with ZIKV showed fetal brain damage in the pups; however, no progression to microcephaly was observed [40].

The introduction of ZIKV to the Americas has been followed by a steady spread of the virus, tied to the range of the arthropod vector which has also increased in recent years [41]. While ZIKV infection rates are certain to rise and fall cyclically, dependent at least in part on weather patterns (particularly rainfall patterns and consequent mosquito density), it is expected that the overall incidence of human infection will increase as more people are exposed to ZIKV via the bites of infected mosquitos. Control methods are currently focused on reduction or elimination of relevant vector populations, including the deployment of genetically modified mosquitos in order to reduce vector populations [42]. In addition, the development and testing of several putative vaccine candidates has also begun [43]. While these techniques probably represent the best-case approach for dealing with ZIKV, the release of genetically modified organisms is a topic that requires intense study and oversight by the FDA before it is approved. Furthermore, the efforts to develop and license a ZIKV vaccine will require at least several more years before such a product could become commercially available. As such, the development and characterization of the major aspects of ZIKV biology, including the neurovirulence and interactions between the immune system and ZIKV, will be required in order to fully support vaccine and drug design. While no animal model may offer a complete, one-stop solution to understanding ZIKV biology, we believe that the *M. domestica* model for studying ZIKV pathogenesis offers unique opportunities to study the effects of CZS in a system that better represents human immunology and pre/post-natal interactions, while allowing for statistically meaningful studies with large numbers of immunocompetent animals.

In summary, using the intra-cerebral route of inoculation, we infected *M. domestica* pups at embryonic and early fetal stages of development and, 74 days later, well beyond the age of weaning (56 days), we observed ZIKV replication and consequent pathogenesis in neuronal tissue. One infected animal exhibited a significant retardation in body and head growth. Its four infected littermates appeared to be anatomically normal, as did the other 11 animals that had been inoculated with ZIKV, as well as all 10 animals inoculated with PBS.

These data suggest that laboratory opossums can be an important new model for studying the effects of ZIKV replication *in vivo* and perhaps also for testing drug therapies, as well as vaccines and other strategies for preventing pathologies caused by ZIKV infection. Moreover, it is possible that ZIKV persists long-term in the brains of some humans after in utero infection, as it does in opossums, without causing any anatomic developmental abnormalities. Some of the infected opossums also did not exhibit any obvious brain pathologies. Some humans who were infected with ZIKV in utero might continue to harbor ZIKV in their brains (an immunologically privileged site). If they do, some of them might develop brain pathologies as some opossums do, and some of them might not develop brain pathologies. The opossum may prove to be a critical model for research on the long-term effects of in utero ZIKV infection during childhood development and into adulthood.

## ACKNOWLEDGEMENTS

We thank Dr. Arthur Porto for directing the work involving head and body measurements, and preparation of heads for photography; Dr. Ana Cristina Leandro for assistance in preparing the photographs of the heads; Dr. Michael Mahaney for advice on statistical analysis; and Alejandro Reyes, Gabriel Lopez, and Cinthya Fuentes-Tapia for expert care of the animals and collection of anatomic data from them.

## REFERENCES

1. Zanluca C, Melo VC, Mosimann AL, Santos GI, Santos CN, Luz K. First report of autochthonous transmission of Zika virus in Brazil. Memórias do Instituto Oswaldo Cruz. 2015 Jun;110(4):569–72.

2. Sarno M, Sacramento GA, Khouri R, do Rosário MS, Costa F, Archanjo G, Santos LA, Nery Jr N, Vasilakis N, Ko AI, de Almeida AR. Zika virus infection and stillbirths: a case of hydrops fetalis, hydranencephaly and fetal demise. PLoS neglected tropical diseases. 2016 Feb 25;10(2):e0004517.

3. Hayes EB. Zika virus outside Africa. Emerging infectious diseases. 2009 Sep;15(9):1347.

4. Dick GW, Kitchen SF, Haddow AJ. Zika virus (I). Isolations and serological specificity. Transactions of the royal society of tropical medicine and hygiene. 1952 Sep 1;46(5):509–20.

5. De Carvalho NS, De Carvalho BF, Fugaça CA, Dóris B, Biscaia ES. Zika virus infection during pregnancy and microcephaly occurrence: a review of literature and Brazilian data. The Brazilian Journal of Infectious Diseases. 2016 May 1;20(3):282–9.

6. Calvet G, Aguiar RS, Melo AS, Sampaio SA, De Filippis I, Fabri A, Araujo ES, de Sequeira PC, de Mendonça MC, de Oliveira L, Tschoeke DA. Detection and sequencing of Zika virus from amniotic fluid of fetuses with microcephaly in Brazil: a case study. The Lancet infectious diseases. 2016 Jun 1;16(6):653–60.

7. Driggers RW, Ho CY, Korhonen EM, Kuivanen S, Jääskeläinen AJ, Smura T, Rosenberg A, Hill DA, DeBiasi RL, Vezina G, Timofeev J. Zika virus infection with prolonged maternal viremia and fetal brain abnormalities. New England Journal of Medicine. 2016 Jun 2;374(22):2142–51.

8. Marrs C, Olson G, Saade G, Hankins G, Wen T, Patel J, Weaver S. Zika virus and pregnancy: a review of the literature and clinical considerations. American journal of perinatology. 2016 Jun;33(07):625–39.

9. Pawitwar SS, Dhar S, Tiwari S, Ojha CR, Lapierre J, Martins K, Rodzinski A, Parira T, Paudel I, Li J, Dutta RK. Overview on the current status of Zika virus pathogenesis and animal related research. Journal of Neuroimmune Pharmacology. 2017 Sep 1;12(3):371–88.

10. Dudley DM, Aliota MT, Mohr EL, Weiler AM, Lehrer-Brey G, Weisgrau KL, Mohns MS, Breitbach ME, Rasheed MN, Newman CM, Gellerup DD. A rhesus macaque model of Asian-lineage Zika virus infection. Nature communications. 2016 Jun 28;7:12204.

11. Mavigner M, Raper J, Kovacs-Balint Z, Gumber S, O’neal JT, Bhaumik SK, Zhang X, Habib J, Mattingly C, McDonald CE, Avanzato V. Postnatal Zika virus infection is associated with persistent abnormalities in brain structure, function, and behavior in infant macaques. Science translational medicine. 2018 Apr 4;10(435):eaao6975.

12. Osuna CE, Lim SY, Deleage C, Griffin BD, Stein D, Schroeder LT, Omange R, Best K, Luo M, Hraber PT, Andersen-Elyard H. Zika viral dynamics and shedding in rhesus and cynomolgus macaques. Nature medicine. 2016 Dec;22(12):1448.

13. Waldorf KM, Stencel-Baerenwald JE, Kapur RP, Studholme C, Boldenow E, Vornhagen J, Baldessari A, Dighe MK, Thiel J, Merillat S, Armistead B. Fetal brain lesions after subcutaneous inoculation of Zika virus in a pregnant nonhuman primate. Nature medicine. 2016 Nov;22(11):1256.

14. Hirsch AJ, Roberts VH, Grigsby PL, Haese N, Schabel MC, Wang X, Lo JO, Liu Z, Kroenke CD, Smith JL, Kelleher M. Zika virus infection in pregnant rhesus macaques causes placental dysfunction and immunopathology. Nature communications. 2018 Jan 17;9(1):263.

15. Osuna CE, Lim SY, Deleage C, Griffin BD, Stein D, Schroeder LT, Omange R, Best K, Luo M, Hraber PT, Andersen-Elyard H. Zika viral dynamics and shedding in rhesus and cynomolgus macaques. Nature medicine. 2016 Dec;22(12):1448.

16. Miner JJ, Sene A, Richner JM, Smith AM, Santeford A, Ban N, Weger-Lucarelli J, Manzella F, Rückert C, Govero J, Noguchi KK. Zika virus infection in mice causes panuveitis with shedding of virus in tears. Cell reports. 2016 Sep 20;16(12):3208–18.

17. Cugola FR, Fernandes IR, Russo FB, Freitas BC, Dias JL, Guimarães KP, Benazzato C, Almeida N, Pignatari GC, Romero S, Polonio CM. The Brazilian Zika virus strain causes birth defects in experimental models. Nature. 2016 Jun;534(7606):267.

18. Li C, Xu D, Ye Q, Hong S, Jiang Y, Liu X, Zhang N, Shi L, Qin CF, Xu Z. Zika virus disrupts neural progenitor development and leads to microcephaly in mice. Cell stem cell. 2016 Jul 7;19(1):120–6.

19. Van den Broek MF, Müller U, Huang S, Aguet M, Zinkernagel RM. Antiviral defense in mice lacking both alpha/beta and gamma interferon receptors. Journal of virology. 1995 Aug 1;69(8):4792–6.

20. Deckard DT. Male-to-male sexual transmission of Zika virus—Texas, January 2016. MMWR. Morbidity and mortality weekly report. 2016;65.

21. Foy BD, Kobylinski KC, Foy JL, Blitvich BJ, da Rosa AT, Haddow AD, Lanciotti RS, Tesh RB. Probable non–vector-borne transmission of Zika virus, Colorado, USA. Emerging infectious diseases. 2011 May;17(5):880.

22. Wang ZE, Reiner SL, Zheng S, Dalton DK, Locksley RM. CD4+ effector cells default to the Th2 pathway in interferon gamma-deficient mice infected with Leishmania major. Journal of Experimental Medicine. 1994 Apr 1;179(4):1367–71.

23. Manangeeswaran M, Ireland DD, Verthelyi D. Zika (PRVABC59) infection is associated with T cell infiltration and neurodegeneration in CNS of immunocompetent neonatal C57Bl/6 mice. PLoS pathogens. 2016 Nov 17;12(11):e1006004.

24. Brasil P, Pereira Jr JP, Moreira ME, Ribeiro Nogueira RM, Damasceno L, Wakimoto M, Rabello RS, Valderramos SG, Halai UA, Salles TS, Zin AA. Zika virus infection in pregnant women in Rio de Janeiro. New England Journal of Medicine. 2016 Dec 15;375(24):2321–34.

25. Rasmussen SA, Jamieson DJ, Honein MA, Petersen LR. Zika virus and birth defects— reviewing the evidence for causality. New England Journal of Medicine. 2016 May 19;374(20):1981–7.

26. Cardoso-Moreira M, Halbert J, Valloton D, Velten B, Chen C, Shao Y, Liechti A, Ascencao K, Rummel C, Ovchinnikova S, Mazin P. Gene expression across mammalian organ development. Nature. 2019 June 26;571 (504-509).

27. Shao Q, Herrlinger S, Yang SL, Lai F, Moore JM, Brindley MA, Chen JF. Zika virus infection disrupts neurovascular development and results in postnatal microcephaly with brain damage. Development. 2016 Nov 15;143(22):4127–36.

28. VandeBerg JL, Williams-Blangero S. The laboratory opossum. The UFAW Handbook on the Care and Management of Laboratory and Other Research Animals. 2010 Jan 19;8:246–61.

29. Samollow PB. The opossum genome: insights and opportunities from an alternative mammal. Genome research. 2008 Aug 1;18(8):1199–215.

30. Wheaton BJ, Noor NM, Dziegielewska KM, Whish S, Saunders NR. Arrested development of the dorsal column following neonatal spinal cord injury in the opossum, Monodelphis domestica. Cell and tissue research. 2015 Mar 1;359(3):699–713.

31. Ritter JM, Martines RB, Zaki SR. Zika Virus: Pathology From the Pandemic. Archives of Pathology & Laboratory Medicine. 2017 January 2017; 141(1): 49–59.

32. European Center for Disease Prevention and Control. Rapid Risk Assessment: Zika virus epidemic in the Americas: potential association with microcephaly and Guillain-Barré syndrome. Stockholm: ECDC 2015 Dec 10.

33. Shan C, Xie X, Muruato AE, Rossi SL, Roundy CM, Azar SR, Yang Y, Tesh RB, Bourne N, Barrett AD, Vasilakis N. An infectious cDNA clone of Zika virus to study viral virulence, mosquito transmission, and antiviral inhibitors. Cell host & microbe. 2016 Jun 8;19(6):891–900.

34. Elong Ngono A, Shresta S. Immune response to dengue and Zika. Annual review of immunology. 2018 Apr 26;36:279–308.

35. Schuler-Faccini L. Possible association between Zika virus infection and microcephaly— Brazil, 2015. MMWR. Morbidity and mortality weekly report. 2016;65.

36. Song H, Qi J, Haywood J, Shi Y, Gao GF. Zika virus NS1 structure reveals diversity of electrostatic surfaces among flaviviruses. Nature structural & molecular biology. 2016 May;23(5):456.

37. Winkler G, Randolph VB, Cleaves GR, Ryan TE, Stollar V. Evidence that the mature form of the flavivirus nonstructural protein NS1 is a dimer. Virology. 1988 Jan 1;162(1):187–96.

38. Young PR, Hilditch PA, Bletchly C, Halloran W. An antigen capture enzyme-linked immunosorbent assay reveals high levels of the dengue virus protein NS1 in the sera of infected patients. Journal of clinical microbiology. 2000 Mar 1;38(3):1053–7..

39. Nguyen SM, Antony KM, Dudley DM, Kohn S, Simmons HA, Wolfe B, Salamat MS, Teixeira LB, Wiepz GJ, Thoong TH, Aliota MT. Highly efficient maternal-fetal Zika virus transmission in pregnant rhesus macaques. PLoS pathogens. 2017 May 25;13(5):e1006378.

40. Lazear HM, Govero J, Smith AM, Platt DJ, Fernandez E, Miner JJ, Diamond MS. A mouse model of Zika virus pathogenesis. Cell host & microbe. 2016 May 11;19(5):720–30.

41. Kraemer MU, Sinka ME, Duda KA, Mylne AQ, Shearer FM, Barker CM, Moore CG, Carvalho RG, Coelho GE, Van Bortel W, Hendrickx G. The global distribution of the arbovirus vectors Aedes aegypti and Ae. albopictus. elife. 2015 Jun 30;4:e08347.

42. Alphey L, Benedict M, Bellini R, Clark GG, Dame DA, Service MW, Dobson SL. Sterile-insect methods for control of mosquito-borne diseases: an analysis. Vector-Borne and Zoonotic Diseases. 2010 Apr 1;10(3):295–311.

43. Poland GA, Kennedy RB, Ovsyannikova IG, Palacios R, Ho PL, Kalil J. Development of vaccines against Zika virus. The Lancet Infectious Diseases. 2018 Jul 1;18(7):e211–9.

